# Effect of dependency and tail behavior on a probability inequality occurring in modeling cognitive processes

**DOI:** 10.1101/2025.11.26.690648

**Authors:** Paria Jahansa, Adele Diederich, Hans Colonius

**Author notes:** Department of Psychology, Carl von Ossietzky Universität Oldenburg,. Contributing authors.

## Abstract

A central idea in modeling performance in cognitive tasks is dynamic competition among processes in separate channels, known as “race model”. This model implies a certain inequality between associated probability distributions under rather general conditions. The inequality represents an important empirical test of the race model, but its usefulness is limited since it requires the assumption of stochastic independence between the channels. Using the stop signal paradigm as reference, we investigate more general forms of stochastic dependence that still imply the inequality using the concepts of copula and heavy-tailed marginal distributions.

## 1 Introduction: race model inequalities

The analysis of reaction times (RTs) is a central part of modeling cognitive processes (Algom, Eidels, Hawkins, Jefferson Townsend, 2015; Colonius Diederich, 2020; Luce, 1986; Townsend Ashby, 1983). It is often based on an important concept, the notion of parallel processing: channels or processes act independently, and the earliest finishing process determines the outcome. For example, in the *redundant signals task*, a person responds whenever a signal appears; sometimes one signal is presented, sometimes two, either simultaneously or with a small delay. The stimuli are either from the same modality, or from different modalities, e.g., visual and auditory. In a prominent parallel processing model for this task, each sensory channel independently races towards a threshold triggering detection. The observed overall response time equals the minimum finishing time of the competing channels. The processing times are considered as random variables, and the model is often referred to as *race model* (Raab, 1962). In a variant of this paradigm, a person is asked to respond to both signals presented. In this *conjunctive task*, observed overall response time becomes the maximum finishing time of both signal processing times (Townsend Wenger, 2004).

Another example for parallel processing is the *stop signal task* : participants are instructed to press a button when a go signal appears. On some trials, a stop signal occurs, telling them to inhibit the response. In the classic race model for this task (Logan Cowan, 1984) the go process starts as soon as the stimulus appears. If a stop signal occurs, a stop process is triggered as a second process running in competition with the go process. Whichever process finishes first wins the race. The observed response time is the time of the go process terminating before the stop signal. If the stop signal is the winner, no response can be registered . This non-observability of the time to process the stop signal presents a particular challenge when it comes to estimating the distributions functions of the random processing times involved (Colonius, 1990; Colonius Diederich, 2023; Matzke, Dolan, Logan, Brown Wagenmakers, 2013; Matzke, Verbruggen Logan, 2018).

An important assumption in parallel processing models is *context independence*: the processing speed of an individual ‘racer’ is the same in the single and redundant conditions^1^. To make this more precise, we write *X* and *Y* for non-negative random variables representing parallel processing times for the two signals with distribution functions *F*_*X*_ (*x*), *F*_*Y*_ (*y*), respectively. In the redundant condition, the corresponding bivariate distribution, *F*_*XY*_ (*x, y*), must also exist. Context independence is then defined as *F*_*XY*_ (*x, ∞*) = *F*_*X*_ (*x*) and *F*_*XY*_ (*∞, y*) = *F*_*Y*_ (*y*) holding for all *x, y*.

Interestingly, even under context independence, it is possible that the processing time distribution for a stimulus in the single-stimulus condition may differ from its conditional distribution in the redundant condition, given it is the winner of the race. Specifically, the following has been shown:

### Proposition 1

(Colonius, Ö zyurt Arndt (2001))

*Let X and Y be two non-negative absolutely continuous and stochastically independent random variables with distribution functions F*_*X*_, *F*_*Y*_, *respectively. Assume F*_*X*_ (0) = *F*_*Y*_ (0) = 0 *and P* (*X < Y*) *>* 0, *Then*

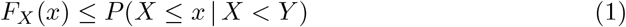

*for all x >* 0.

This *race model inequality* can be interpreted as follows: given that processing time *X* is faster than *Y*, the probability of *X* of staying below a given time *x* is equal or larger than without that condition. Before we proceed, we consider a corresponding result for the conjunctive task (maximum finishing time) that can be obtained from inequality (1):

### Proposition 2

*Let W and Z be two stochastically independent non-negative continuous random variables with distribution functions F*_*W*_, *F*_*Z*_, *respectively. Assume F*_*W*_ (0) = *F*_*Z*_(0) = 0 *and P* (*Z < W*) *>* 0. *Then*

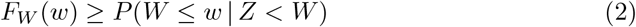

*for all w >* 0.

A proof is in the appendix. In analogy to race model inequality (1), the interpretation of 2 is as follows: given that processing time *W* is longer than *Z*, the probability of *W* of staying below a given time *w* is equal or smaller than without that condition.

Both inequalities appear plausible, and they present a strong test for a given race model. The proof of the propositions was critically based on assuming stochastic independence (SI) among the random processing times in the model. As it turns out, however, SI is not a necessary assumption; indeed, investigating race models for the stop signal task, Colonius, Jahansa, Joe Diederich (2024) present parametric examples with stochastic *dependency* such that the inequalities are still satisfied for specific parametric settings. Thus, the ultimate goal is to characterize exactly those race models that predict inequalities (1) or (2) but, according to our knowledge, only partial results exist up to now. The focus here is to investigate (i) the effect of the type of stochastic dependency among the “racers” and (ii) the effect of some properties of the marginal distributions, like tail behavior, on the inequalities.

The next section sets the stage by presenting a sufficient condition for inequality 1 derived in Colonius et al. (2024); this condition will be the main reference of our study. It will be illustrated by a bivariate normal distribution example in the next section. The subsequent section introduces three relevant statistical concepts: (i) the *copula* allowing one to study the type of stochastic dependency separately from specific marginal distributions, (ii) *heavy-tailed distributions*, and (iii) *bivariate tail-dependence*. Three specific copula types covering different tail dependencies and two marginal distributions with different tail heaviness are introduced. Using the stop signal race model as a standard reference, section 5 combines different copulas and marginals to investigate when the sufficient conditions for inequality 1 and 2 are satisfied.

Given that no theoretical results exist that completely characterize the race model inequalities, our exposition remains largely exploratory. Nonetheless, the numerical analyses of a finite number of well-known copulas and marginals should be useful for further steps toward the ultimate goal.

## 2 Sufficient conditions and a counterexample

We start by referring to the announced result providing sufficient conditions for inequality (1), or it reverse, to hold:

### Proposition 3

(Colonius et al. (2024))

*Let X and Y be random variables with a joint density and conditional (cumulative) distribution function F*_*Y* |*X*_ (*y*|*x*).

a. *If g*(*z*) = 1 *− F*_*Y* |*X*_ (*z*|*z*) *is decreasing in z, then*

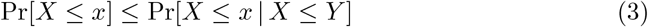

*for all x*.
b. *If g*(*z*) = 1*−F*_*Y* |*X*_ (*z*|*z*) *is* increasing *in z, then the inequality reverses such that*

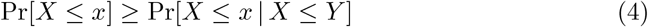

*for all x*.

(Note: “decreasing” is defined as non-increasing, “increasing” as non-decreasing).

The following example from Colonius et al. (2024) illustrates the proposition: depending on the parameter setting one finds the inequality, or its converse, satisfied.

### Example 1

*Let* (*X, Y*)^*′*^ *be a bivariate normal random vector:*

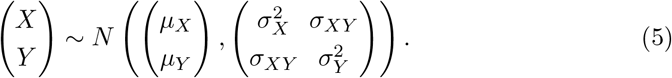

*Then, for the conditional distribution*,

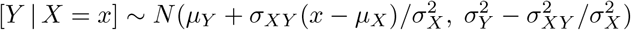

*and, with* Φ *denoting the standard normal distribution function*,

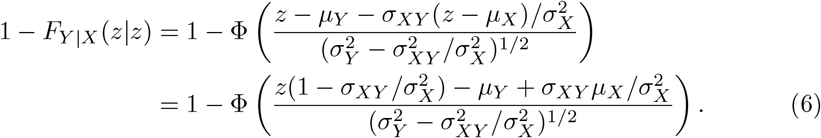

*This is always decreasing in z if* 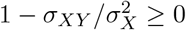

*One can have* 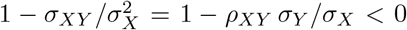 *for ρ*_*XY*_ *positive and σ*_*Y*_ */σ*_*X*_ *sufficiently large. An extreme case is as follows: let ρ*_*XY*_ = 1, *µ*_*X*_ = *µ*_*Y*_ = 0, 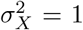, *and Y* = *aX with a >* 1. *Then σ*_*Y*_ = *aσ*_*X*_ = *a and* 1 *− σ*_*XY*_ *σ*_*Y*_ */σ*_*X*_ = 1 *− a <* 0. *The event {X ≤ Y} corresponds to {X ≥* 0*} and*

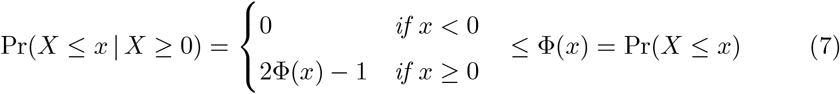

*violating the inequality for all x*.

This example suggests that the (non-)violation of inequality (1) is a function of the type of stochastic dependency between the random variables. For a more systematic investigation, the next section introduces some relevant statistical tools.

## 3 Copulas and tail behavior

### 3.1 Copulas

A *copula* is defined as a function that specifies how a bivariate (in general, multivariate) distribution is related to its one-dimensional marginal distributions. It allows one to assess stochastic dependency between the two random variables separately from the choice of the marginal distributions. Thus, copulas are a natural tool to investigate which combination of stochastic dependency and marginal distributions will lead to a (non-)violation of the inequalities. One formal definition is

#### Definition 1

*A 2-dimensional copula is a 2-dimensional cumulative distribution function C defined on the unit square* [0, 1]^2^ *with uniformly distributed marginals on* [0, 1],

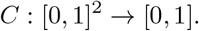

*For any u, v ∈* [0, 1], *C*(*u, v*) *lies in* [0, 1].

We only consider continuous, two-dimensional (*d* = 2) copulas here; for the general case, we refer to Czado (2019); Joe (2014); Nelsen (2006), and Durante Sempi (2016). An important result follows directly from the celebrated *Sklar’s theorem*, a simple version of which is the following:

#### Proposition 4

(Sklar (1959))

a. *For a bivariate distribution F*_*XY*_ *with margins F*_*X*_ *and F*_*Y*_, *the copula C associated with F*_*XY*_ *exists with uniform margins on* [0, 1]

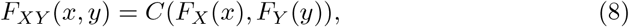

*for* (*x, y*) *∈* **R**^2^.
b. *If F*_*XY*_ *is a bivariate distribution function with continuous margins F*_*X*_, *F*_*Y*_ *and quantile functions (inverses)* 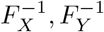, *then*

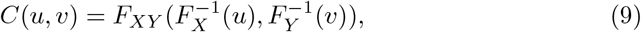

*for* (*u, v*) ∈ [0, 1]^2^.

Here is an example of a copula example that will further be considered below:

#### Example 2 (Gumbel copula)

*The bivariate Gumbel copula with parameter θ is defined as*

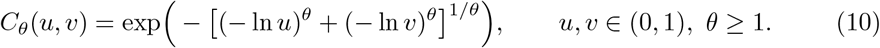

The next remark lists several relevant definitions and properties of copula theory; details and proofs can be found in the references given above. Let (*X, Y*) be a pair of random variables with distribution function *F*_*XY*_, univariate distribution functions *F*_*X*_ and *F*_*Y*_, and copula *C*.

#### Remark 1

survival copula:

i. *with univariate survival functions* 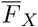 *and* 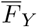, *the joint survival function is given by*

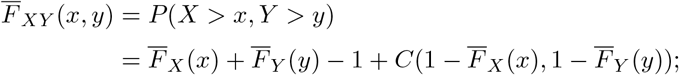

*then, define function Ĉ from* [0, 1]^2^ *into* [0, 1],

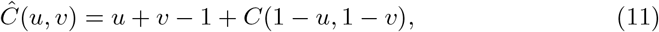

*so that*

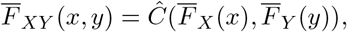

*with Ĉ being a copula, the survival copula*.
ii. copula density: *for an absolutely continuous copula C*(*u, v*) *copula, density is defined as*

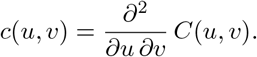

*Then, for a pair random variables* (*X, Y*) *with density f*_*XY*_ (*x, y*),

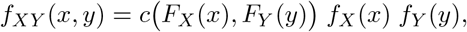

*and the conditional density equals*

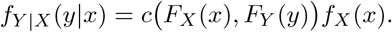

*Conditioning on Y is analogous*.
iii. Conditional copula *(defined as conditional distribution of V given U = u)*

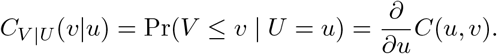
iv. *for a pair of random variables* (*X, Y*), *the* conditional distribution of *Y* given *X* = *x*,

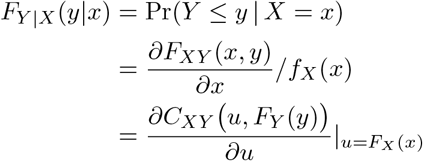

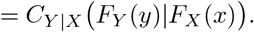

These properties are illustrated next by adding marginals to the Gumbel copula example.

#### Example 3 (Gumbel copula with exponential marginals)

*The bivariate Gumbel copula with parameter θ is defined as*

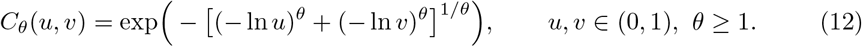

*Inserting exponential marginals*,

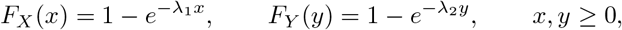

*where λ*_1_, *λ*_2_ *>* 0 *are rate parameters, this becomes:*

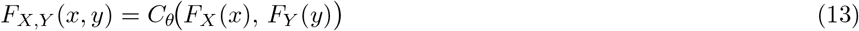

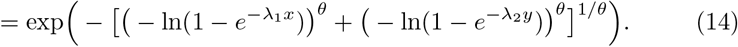

*The corresponding joint probability density function is*

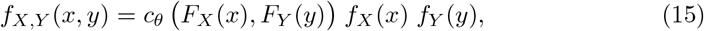

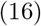

*where*

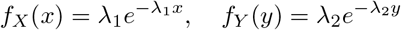

*and*

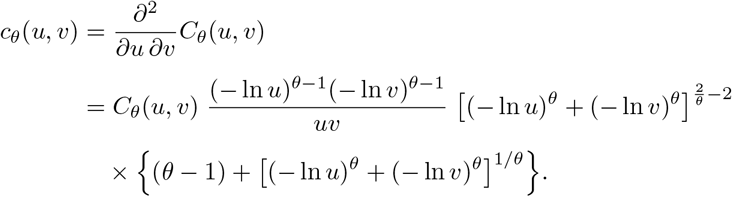

*The conditional copula of* (*V* |*U* = *u*) *is*

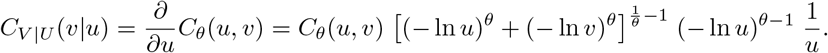

### 3.2 Tail behavior

Tail behavior refers to properties of distributions for very large values (including infinitely large values) at the right or left end. A single distribution may be heavy-tailed, while a pair of distributions may expose specific tail dependency. We consider both cases separately.

#### 3.2.1 Heavy-tailed distributions

Simply, heavy tail of a distribution means that there is a larger probability of getting very large values; however, different concepts exist in the literature. For example, a Gaussian distribution is said to have a heavier tail than a uniform, an exponential a heavier tail than a Gaussian. A key feature of the definition is the comparison distribution chosen (for a recent reference, see Nair, Wierman Zwart (2022)).

##### Definition 2

*A distribution function F is said to be* heavy-tailed *if and only if for all µ >* 0

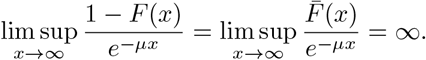

*Otherwise, F is light-tailed. A random variable X is said to be heavy-tailed (light-tailed) if its distribution function is heavy-tailed (light-tailed)*.

Thus, this definition uses the exponential as comparison point to define the class of heavy-tailed distributions (the exponential itself is light-tailed). Different distribution families exhibit different tail behaviors. For example, the normal distribution is known to have light tails, implying a low probability of extreme values. In contrast, the *t*-distribution has heavier tails compared to the normal, and the Cauchy distribution exhibits even heavier tails, assigning relatively high probability to extreme observations.

#### 3.2.2 Bivariate tail dependence

Tail dependence is a measure of strength of dependence in the joint lower or joint upper tail of a multivariate distribution; here, we only consider the bivariate case.

##### Definition 3

*Let X and Y be continuous random variables with distribution functions F*_*X*_ *and F*_*Y*_, *respectively. The* upper bivariate tail dependence coefficient *λ*_*U*_ *is the limit (if it exists)*

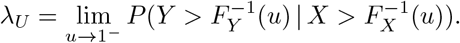

*Similarly, the* lower bivariate tail dependence coefficient *λ*_*L*_ *is the limit (if it exists)*

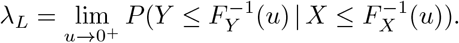

A definition in terms of bivariate copulas is the following:

##### Definition 4

*A bivariate copula C has* upper tail dependence *if*

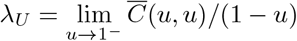

*exists with λ*_*U*_ *∈* (0, 1], *and* no upper tail dependence *if λ*_*U*_ = 0. *Similarly, C has* lower tail dependence *if*

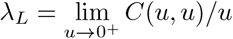

*exists with λ*_*L*_ *∈* (0, 1], *and* no lower tail dependence *if λ*_*L*_ = 0.

Thus, the upper tail dependence coefficient indicates the asymptotic limit of the probability that one random variable exceeds a high quantile, given that the other random variable exceeds a high quantile. Similar interpretation holds for the lower tail dependence coefficient. There are various types of copulassuch as the Gaussian, Gumbel, and Claytoneach capturing different forms of dependence. For instance, the Gaussian copula is limited to modeling linear positive and negative dependencies and lacks tail dependence, whereas the Gumbel and Clayton copulas can capture upper and lower tail dependencies, respectively (Czado, 2019; Joe, 2014; Nelsen, 2006).

## 4 Investigating the race model inequalities in the stop signal model

The stop signal task mentioned in the introduction is used to study cognitive control and response inhibition (Verbruggen et al., 2019). The race model (Logan Cowan, 1984) describes performance in this task as a race between two random variables: go processing time, *T*_*go*_, and stop processing time, *T*_*stop*_. The model is based on two main assumptions: *context independence*, meaning the distribution of go processing time remains the same whether or not a stop signal is presented, and *stochastic independence*, meaning the two random variables, *T*_*go*_ and *T*_*stop*_, are stochastically independent.

Under these assumptions, the model obeys race model inequality (1) if

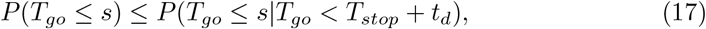

which is now written in terms of random variables in the race model, *T*_*go*_ and *T*_*stop*_, and where *t*_*d*_ denotes a constant, the time the stop signal is delayed relative to the go signal (SSD) in a given trial. As pointed out in the introduction, the present study started from the empirical observation that, sometimes, deviations from this model prediction occur (Colonius et al., 2001). In particular, Bissett, Jones, Poldrack Logan (2021) observed that severe violations occur primarily at shorter SSD values, interpreting this as a failure of the context independence assumption. Alternatively, in a previous paper (Colonius et al., 2024), we suggested that such deviations can be explained by an effect of stochastic dependency between the “racers” in the model. Specifically, it was shown that the inequality is always satisfied for Gumbels bivariate exponential distribution, which is characterized by negative dependence only. In contrast, a violation of the inequality was observed for a correlated ex-Gaussian distribution.

Further progress in resolving this problem for stop signal modeling hinges upon finding conditions under which inequality (17), or its reversal, are satisfied. As a first step, we focus on a condition that was found to be sufficient for the inequality to hold, as stated in Proposition 3:

For random variables *X* and *Y*, conditional (cumulative) distribution function *F*_*Y* |*X*_ (*y*|*x*),

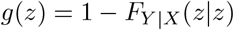

is decreasing in *z*. In the notation of the stop signal model, this equals

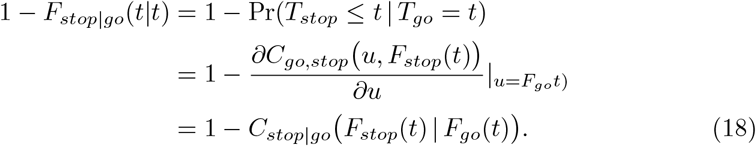

### 4.1 Choosing marginals and copulas

The task is to check whether function (18) is monotonic, either non-decreasing or non-increasing, for a given choice of a copula and corresponding marginals distributions. Proposition 3 then guarantees that inequality (17), or its reversal, is satisfied. Obviously, there is an infinite number of copulas and marginals to choose from. Here we settle on three copula types that exhibit different tail dependence: Gumbel and Survival Gumbel copula capture upper and lower tail dependence, respectively, and the Gaussian copula lacks tail dependence. Moreover, we select two types of marginals, the exponential and the log-normal.

#### 4.1.1 Marginals

The different exponential and the log-normal are both are right-skewed, which is consistent with the typical shape of reaction time distributions. The exponential distribution is light-tailed, while the log-normal distribution exhibits a heavier tail. The cumulative distribution function of

- Exponential distribution with parameter *λ >* 0 is:

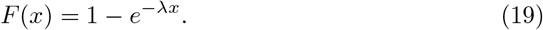
- Log–normal distribution with parameters *µ ∈* **R** and *σ >* 0 is:

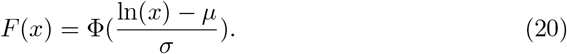

with Φ denoting the standard normal cumulative distribution.

#### 4.1.2 Gumbel copula

This copula was given in equation (13) with exponential marginals in example 2. Considering exponential and lognormal marginals *u* = *F*_*X*_ (*x*) = 1 *− e*^*−λx*^ and 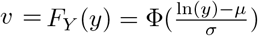 and using (12), the joint distribution is

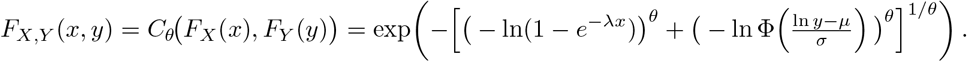

#### 4.1.3 Survival Gumbel copula

The copula specified in (11), together with exponential and lognormal marginals *F*_*X*_ (*x*) = 1 *− e*^*−λx*^ and 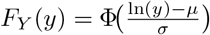, yields the joint distribution

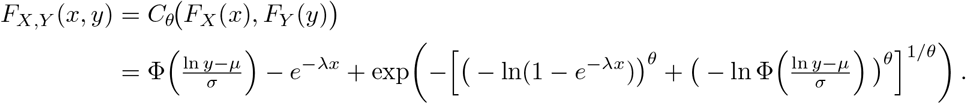

#### 4.1.4 Gaussian copula

For a given correlation matrix *ρ ∈* [*−*1, 1]^2*×*2^, the Gaussian copula with parameter *ρ* is

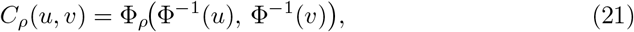

where Φ^*−*1^ is the inverse cumulative distribution function of a standard normal and Φ_*ρ*_ is the joint CDF of a bivariate normal distribution with mean vector 0 and correlation matrix *ρ*. The joint distribution of two random variables *X* and *Y* with exponential and lognormal marginals *F*_*X*_ (*x*) = 1 *− e*^*−λx*^ and 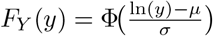, and a Gaussian copula is

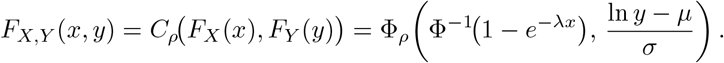

### 4.2 Conditional copulas

In order to investigate the sufficient condition, we need to insert the specific conditional copulas to into equation 18 (derivations are found in appendix B).

Let *u* = *F*_*go*_(*t*), and *v* = *F*_*stop*_(*t*). For any *t > t*_*d*_, the conditional copula *C*_*stop*|*go*_ of

- Gumbel copula with parameter *θ ≥* 1 is:

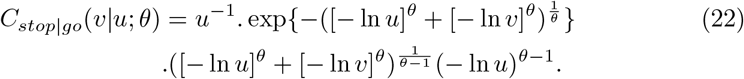
- Survival Gumbel copula with parameter *θ ≥* 1 is defined as the reflected Gumbel copula:

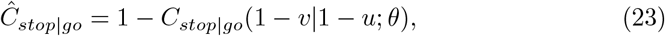

where *C*_*stop*|*go*_ denotes the conditional Gumbel copula.
- Gaussian Copula with parameter *−*1 *< ρ <* 1 is:

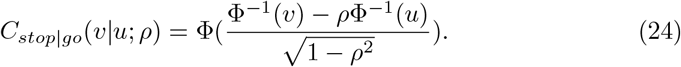

## 5 Numerical results

The first step in the numerical analysis involves selecting parameters for each distribution and copula function. The parameters of the marginal distributions were chosen so that the mean reaction times fall within the range of approximately 150 to 300 ms. Copula parameters were selected to cover a variety of dependency structures, including different levels of overall dependence and tail dependence (both heavy and light). For the Gumbel and survival Gumbel copulas, two different values of the dependence parameter were considered: *θ ∈ {*1.2, 5*}*. For the Gaussian copula, the correlation parameter (copula parameter) was chosen as *ρ* ∈ {−0.8, 0.9} to represent strong negative and strong positive dependence, respectively. Among the eligible parameter values for the exponential distribution, *λ* = 0.005 yields a mean around 200 ms. For the log-normal distribution, choosing *µ* = 1 and 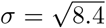 also results in a mean near 200 ms. The numerical calculations were also performed for three different SSDs, *t*_*d*_ ∈ {5, 60, 110}, in order to probe the effect of SSD on the results.

The results are presented separately for each copula. We compare the effects of different marginal distributions alongside the dependency structure of each copula.

### 5.1 Gumbel copula

Figure 1 illustrates 1 *− C*_*stop*|*go*_ as a function of *t* for the Gumbel copula with parameter *θ* = 1.2, corresponding to Kendall’s *τ* = 0.16, upper tail dependence *λ*_*U*_ = 0.21, and no lower tail dependence (*λ*_*L*_ = 0). This represents a case of weak upper tail dependence. Panels (A) and (B) show cases where *T*_*stop*_ has a heavier tail and *T*_*go*_ has a heavier tail, respectively. In panel (B), 1 *− C*_*stop*|*go*_ decreases with *t* indicating no violation of the inequality (17). In contrast, panel (A) shows that 1 *− C*_*stop*|*go*_ is non-monotonic in *t* where *T*_*stop*_ has the heavier tail, suggesting the possibility of a violation .

**Fig. 1:**
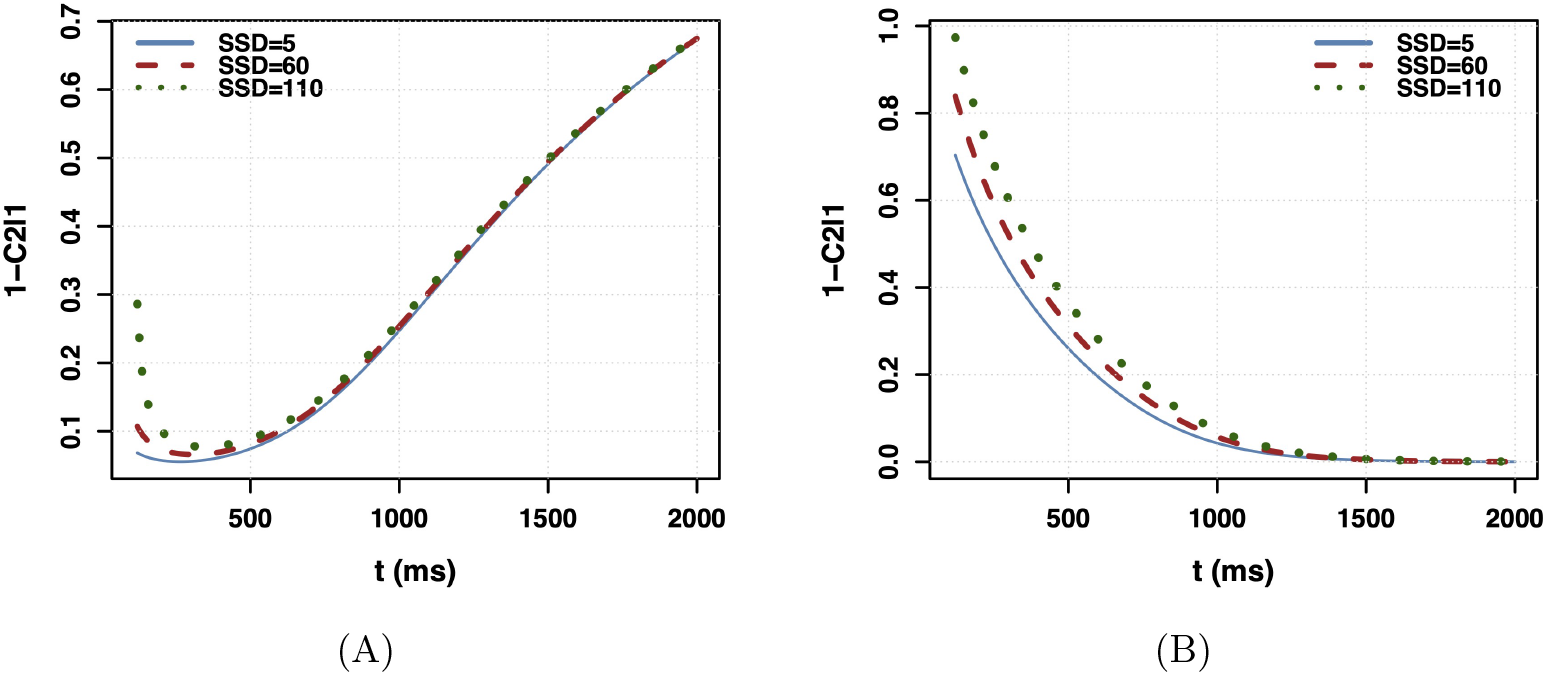
Behavior of 1 *− C*_stop|go_(*t*) for the Gumbel copula with parameter *θ* = 1.2. Panel (A): *T*_go_ *∼* Exp(0.005), *T*_stop_ *∼* Lognormal(1,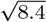). Here, the stop process has the heavier tail, and 1 *− C*_stop|go_(*t*) is non-monotonic in *t*. Panel (B): *T*_go_ *∼* Lognormal(1, 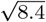), *T*_stop_ *∼* Exp(0.005). In this case, the go process has the heavier tail, and 1 *− C*_stop|go_(*t*) is decreasing in *t*.

Figure 2 is similar to Figure 1, but with a higher Gumbel copula parameter, *θ* = 5, corresponding to Kendall’s *τ* = 0.8, upper tail dependence *λ*_*U*_ = 0.85, indicating strong upper tail dependence. In this case, panel (A) shows an increasing trend of 1 *− C*_*stop*|*go*_ across all *t* when *T*_*stop*_ has the heavier tail, indicating a severe violation of the sufficient conditionsuch that the reverse of inequality (17) is satisfied. In contrast, panel (B), where *T*_*go*_ has a heavier tail, continues to show no violation, with 1*− C*_*stop*|*go*_ decreasing in *t*.

**Fig. 2:**
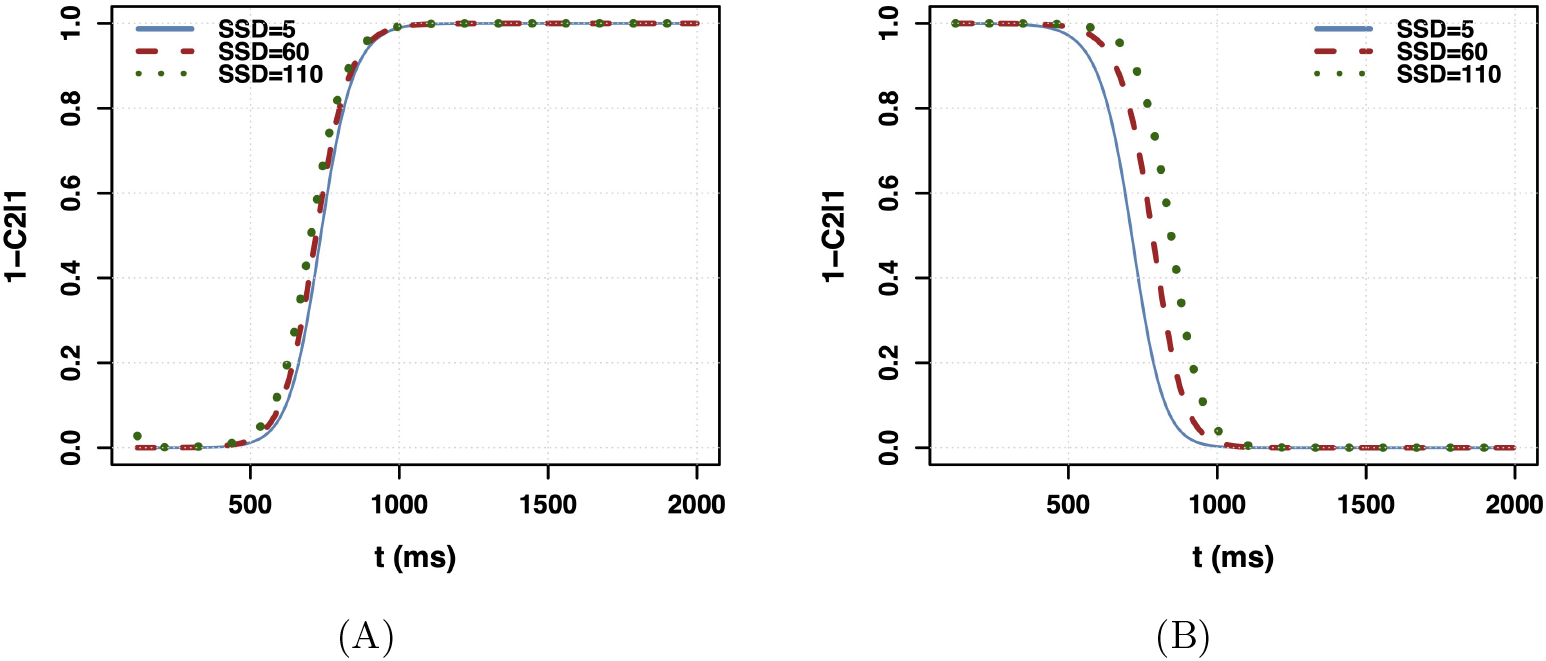
Behavior of 1 *− C*_stop|go_(*t*) for the Gumbel copula with parameter *θ* = 5. Panel (A): *T*_go_ *∼* Exp(0.005), *T*_stop_ *∼* Lognormal(1, 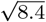). Here, the stop process has the heavier tail, and 1 *− C*_stop|go_(*t*) is increasing in *t*. Panel (B): *T*_go_ *∼* Lognormal(1, 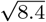), *T*_stop_ *∼* Exp(0.005). In this case, the go process has the heavier tail, and 1 *− C*_stop|go_(*t*) is decreasing in *t*.

### 5.2 Survival Gumbel copula

Figures 3 and 4 illustrate 1 *− C*_*stop*|*go*_ for the Survival Gumbel copula with the parameters *θ* = 1.2 and *θ* = 5, respectively. The survival Gumbel copula exhibits lower tail dependence, reflecting the upper tail dependence of the Gumbel copula. For *θ* = 1.2, the copula has Kendall’s *τ* = 0.16, lower tail dependence *λ*_*L*_ = 0.21, and no upper tail dependence (*λ*_*U*_ = 0).

**Fig. 3:**
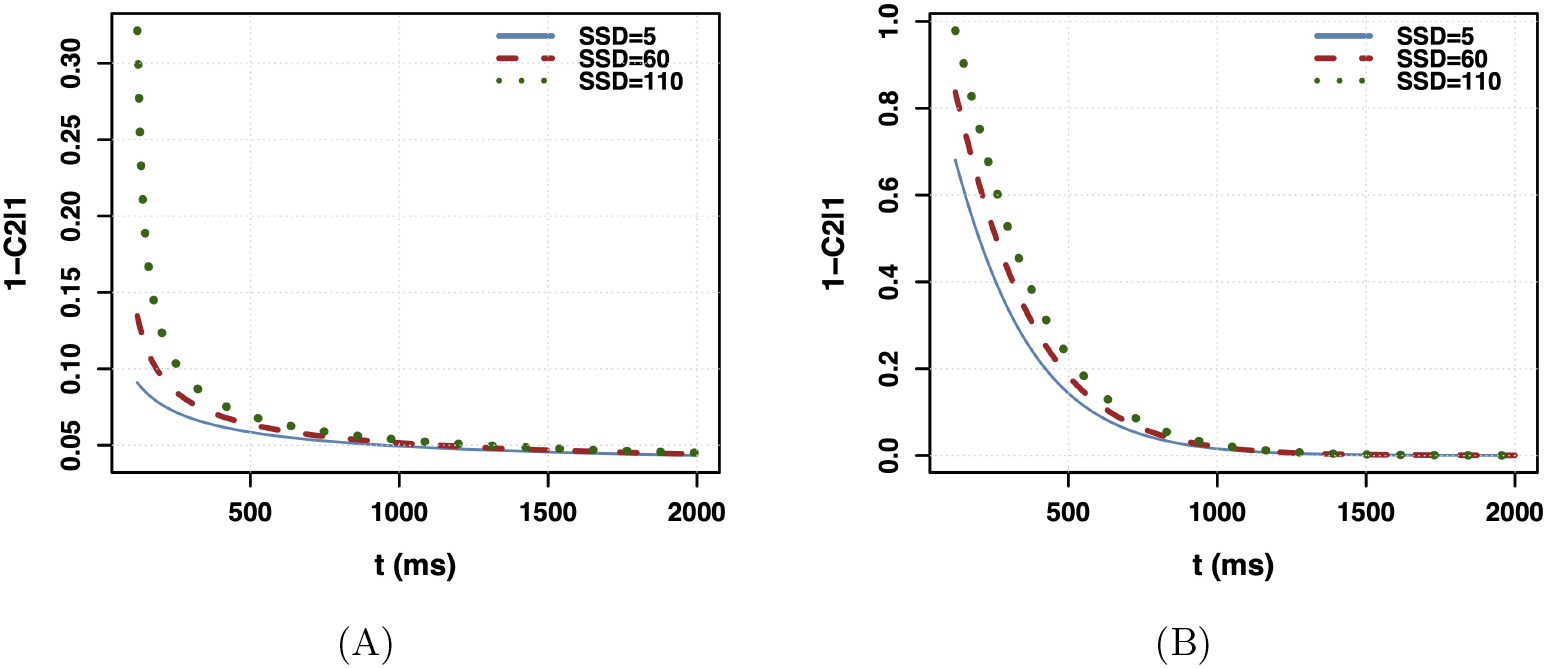
Behavior of 1 *− C*_stop|go_(*t*) for the Survival Gumbel copula with parameter *θ* = 1.2. Panel (A): *T*_go_ *∼* Exp(0.005), *T*_stop_ *∼* Lognormal(1, 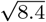). Here, the stop process has the heavier tail, and 1 *− C*_stop|go_(*t*) is decreasing in *t*. Panel (B): *T*_go_ *∼* Lognormal(1, 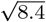), *T*_stop_ *∼* Exp(0.005). In this case, the go process has the heavier tail, and 1 *− C*_stop|go_(*t*) is decreasing in *t*.

**Fig. 4:**
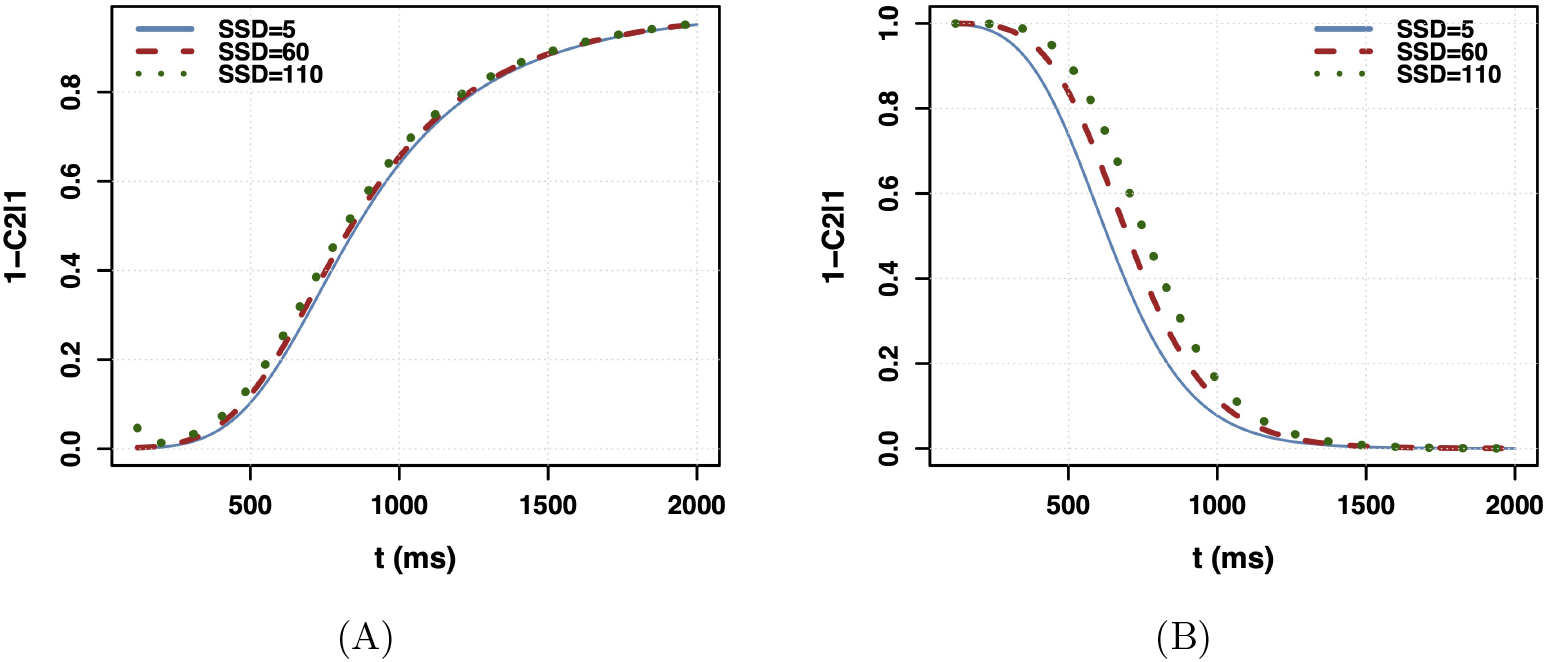
Behavior of 1 *− C*_stop|go_(*t*) for the Survival Gumbel copula with parameter *θ* = 5. Panel (A): *T*_go_ *∼* Exp(0.005), *T*_stop_ *∼* Lognormal(1, 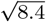). Here, the stop process has the heavier tail, and 1 *− C*_stop|go_(*t*) is increasing in *t*. Panel (B): *T*_go_ *∼* Lognormal(1, 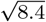), *T*_stop_ *∼* Exp(0.005). In this case, the go process has the heavier tail, and 1 *− C*_stop|go_(*t*) is decreasing in *t*.

Figure 3 shows that, under this weak lower-tail dependence, inequality (17) is satisfied regardless of whether *T*_*stop*_ is heavy-tailed or light-tailed. In contrast, Figure 4 represents much stronger lower-tail dependence with Kendall’s *τ* = 0.8 and *λ*_*L*_ = 0.85. Here, when *T*_*stop*_ is heavy-tailed (panel (A)), inequality (17) is reversed.

### 5.3 Gaussian copula

Figure 5 shows 1 *− C*_*stop*|*go*_ for the Gaussian copula with strong negative dependence (Kendall’s *τ* = *−*0.59). As seen in both panels, 1 *− C*_*stop*|*go*_ decreases with *t* regardless of the marginal distributions of *T*_*stop*_ and *T*_*go*_ implying inequality (17). When the dependence becomes strongly positive, as in Figure 6 (*τ* = 0.71), the behavior changes.

**Fig. 5:**
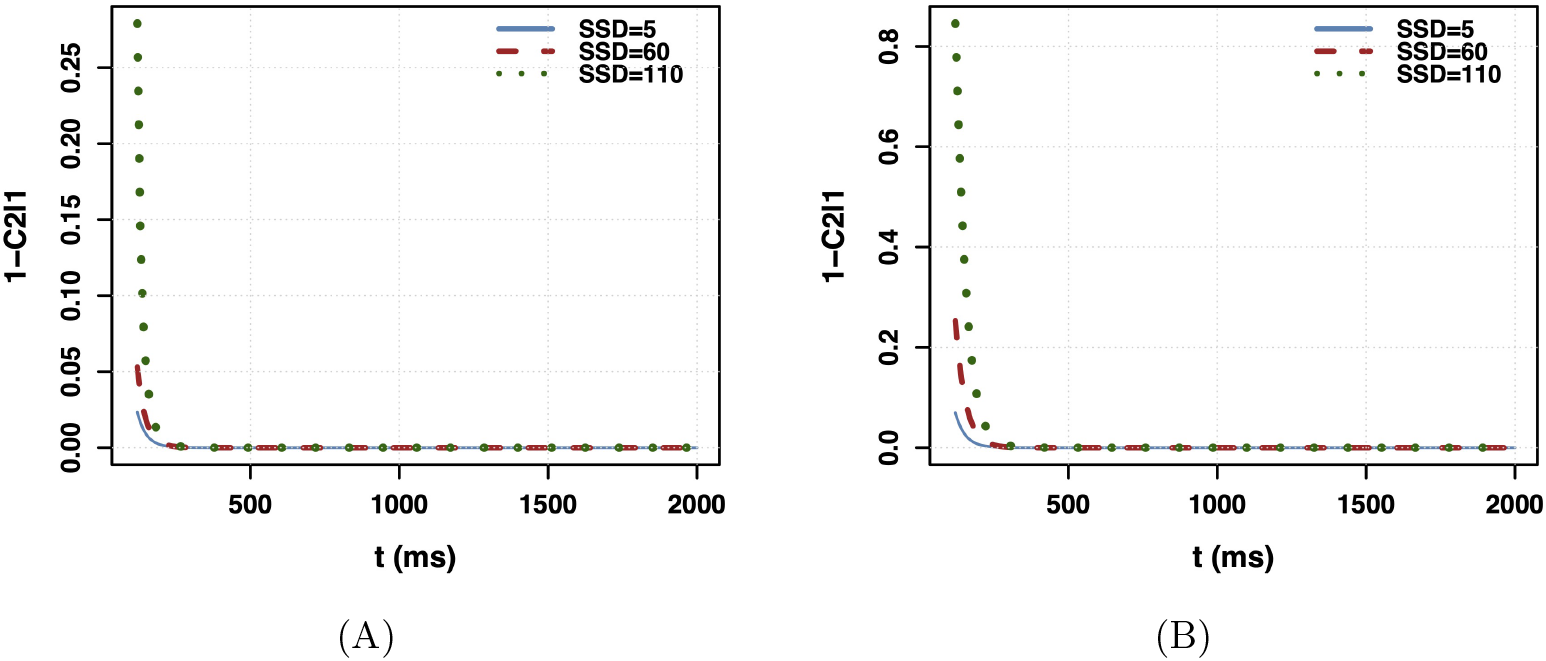
Behavior of 1 *− C*_stop|go_(*t*) for the Gaussian copula with parameter *ρ* = *−*0.8. Panel (A): *T*_go_ *∼* Exp(0.005), *T*_stop_ *∼* Lognormal(1, 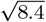). Here, the stop process has the heavier tail, and 1 *− C*_stop|go_(*t*) is decreasing in *t*. Panel (B): *T*_go_ *∼* Lognormal(1, 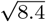), *T*_stop_ *∼* Exp(0.005). In this case, the go process has the heavier tail, and 1 *− C*_stop|go_(*t*) is decreasing in *t*.

**Fig. 6:**
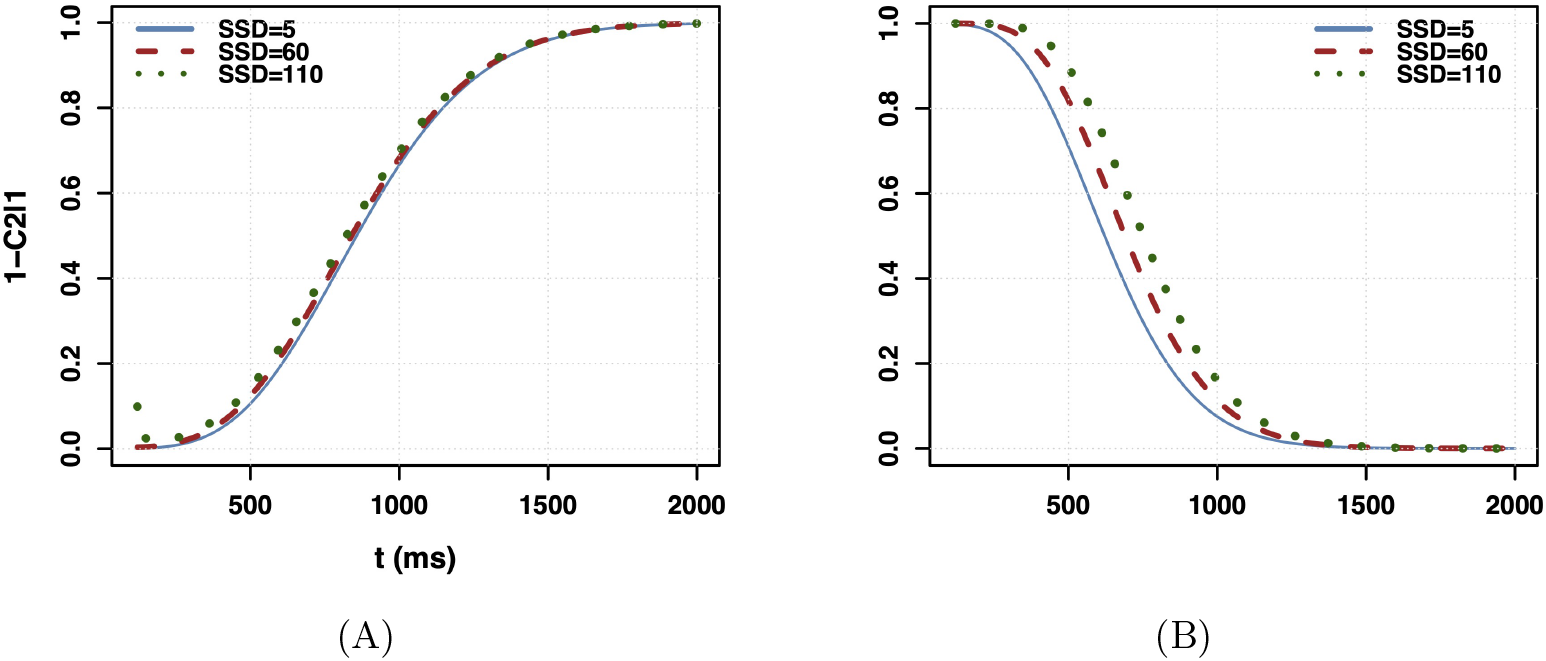
Behavior of 1 *− C*_stop|go_(*t*) for the Gaussian copula with parameter *ρ* = 0.9. Panel (A): *T*_go_ *∼* Exp(0.005), *T*_stop_ *∼* Lognormal(1, 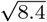). Here, the stop process has the heavier tail, and 1 *− C*_stop|go_(*t*) is increasing in *t*. Panel (B): *T*_go_ *∼* Lognormal(1, 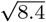), *T*_stop_ *∼* Exp(0.005). In this case, the go process has the heavier tail, and 1 *− C*_stop|go_(*t*) is decreasing in *t*.

In panel (A), where *T*_*stop*_ has a heavier tail than *T*_*go*_, 1 *− C*_*stop*|*go*_ increases with *t*, resulting in a reversal of inequality (17). In contrast, panel (B), where *T*_*go*_ has the heavier tail, shows that the inequality holds.

### 5.4 Summary of numerical results

These numerical findings, demonstrated for the stop signal race model, indicate that violations of probability inequality 1 are closely tied to both the tail behavior of the marginal distributions and the nature of the dependency structure. Specifically, when *T*_*go*_ exhibits a heavier tail, the inequality consistently holds, regardless of the dependence structure. In contrast, if *T*_*stop*_ has a heavier tail, violations tend to occur. These violations are most evident under upper tail dependence between *T*_*go*_ and *T*_*stop*_, where the reverse of the inequality holds. Even in cases of lower tail dependence or independence, the reverse inequality can still arise, particularly under strong positive dependence. This interplay between marginal tail behavior and copula structure highlights the complex conditions under which the inequality may fail or reverse. In essence, the likelihood of a violation increases when the variable associated with stopping behavior dominates in the tail and aligns with a dependence structure that amplifies joint extreme outcomes.

Therefore, both the heaviness of the tails and the type of dependence must be considered together to understand and predict when the inequality may not hold.

## 6 Summary and conclusion

The race model inequality and its reverse, equations 1 and 2, provide an important empirical test of any stochastic race model. Therefore, it is crucial to determine the exact conditions under which these inequalities are valid but, to the best of our knowledge, only partial, sufficient solutions exist in the literature. In the context of the stop signal paradigm, it had been shown previously (Colonius et al., 2001) that stochastic independence between the “racers” suffices for inequality 1 to hold, and Colonius et al. (2024) presented examples where the inequalities remain valid even under stochastic dependency. Based on the sufficient condition stated in proposition 3, here we undertake a further step by considering how validity of the inequalities is related to (i) the type of tail-dependency of the copula determining stochastic dependence and (ii) the tail-heaviness of the marginal distributions and the interaction between the two.

Importantly, our results are limited because they are based on a sufficient condition (3) for the race model inequality; thus, finding a non-monotonic function *g*(*z*) = 1 *− F*_*Y* |*X*_ (*z*|*z*) (see proposition 3) may only indicate that the sufficient condition is too strong rather than presenting unambiguous evidence against the inequality. Future work should probe whether –under copula/parameter settings showing non-monotonic *g*(*z*)– the race model inequality is violated or not.

Since it is computationally intensive to examine all types of copulas, three representative copulaseach capturing a distinct form of dependence were selected for numerical analysis. In addition, two different marginal distributions with different tail-heaviness were chosen. The numerical analysis clearly indicated that higher levels of dependence and greater differences in tail heaviness between the “racers” tend to lead to more extreme violations. However, a more comprehensive set of copulas and potential marginals is certainly called for. Moreover, our numerical analyses had to be based on specific parameter choices, and it remains to identify the exact thresholds at which such violations begin to occur. A promising direction for future work would be to analytically determine the parameter range within which violations of the inequalities are likely to occur. This task can become mathematically challenging, especially for copulas with complex functional forms, and could require advanced analytical or numerical techniques. In some cases, it may difficult, or even infeasible, to derive closed-form solutions.

## 7 Appendix

### 7.1 Appendix (A) Proof of Proposition 2

#### Proof 1

*We set W* = 1*/X, Z* = 1*/Y, and w* = 1*/x; then*,

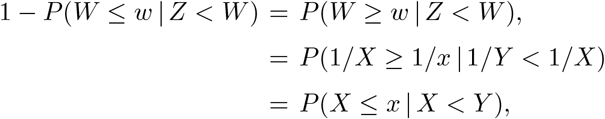

*after multiplying the inequality* 1*/X ≥* 1*/x by both X and x; moreover*,

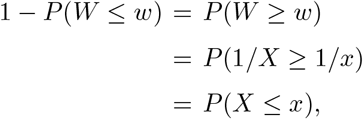

*repeating the multiplication. From (1)*,

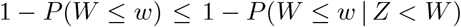

*or*,

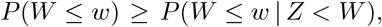

*which was to be shown. The settings are valid because the random variables take positive values almost surely*.

### 7.2 Appendix (B) Partial Derivatives of Copulas

#### 7.2.1 Gumbel

For the Gumbel copula (12), the first partial derivative with respect to *u* is

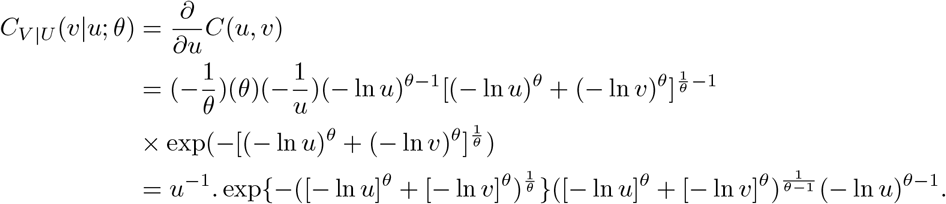

#### 7.2.2 Survival Gumbel

The first partial derivative with respect to *u* of the Survival Gumbel copula (11) is

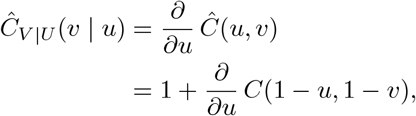

where *C*(1 *− u*, 1 *− v*) is the (standard) Gumbel copula. Since 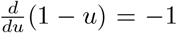, the derivation yields

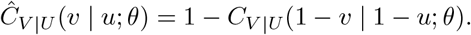

#### 7.2.3 Gaussian

From (21), let *a* = Φ^*−*1^(*u*) and *b* = Φ^*−*1^(*v*). Then

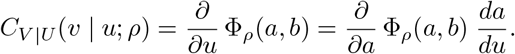

Using the representation of the bivariate standard normal CDF,

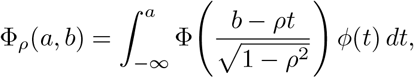

and applying Leibnizs rule, we obtain

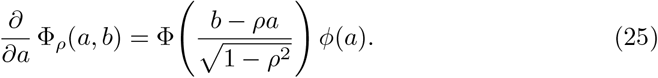

Since Φ(Φ^*−*1^(*u*)) = *u*, differentiating both sides with respect to *u* gives

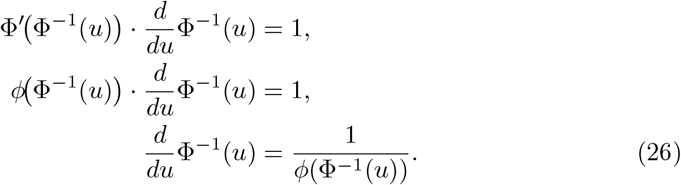

Combining (25) and (26), the conditional copula is

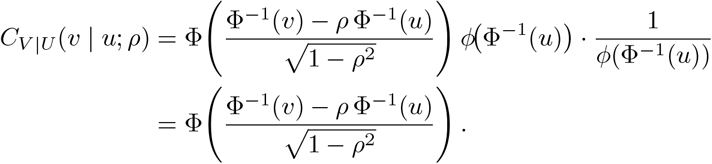

## Footnotes

1 As argued in Miller (2016), without this assumption, the models become unfalsifiable.

## References

Algom, D., Eidels, A., Hawkins, R., Jefferson, B., Townsend, J. (2015). Features of response times: identification of cognitive mechanisms through mathematical modeling. J. Busemeyer, Z. Wang, J. Townsend, & A. Eidels (Eds.), The Oxford Handbook of Mathematical and Computational Psychology (pp. 63–98). New York, NY 10016: Oxford University Press.

Bissett, P.G., Jones, H.M., Poldrack, R.A., Logan, G.D. (2021). Severe violations of independence in response inhibition tasks. Science advances, 7 (12), eabf4355,

Colonius, H. (1990). A note on the stop-signal paradigm, or how to observe the unobservable. Psychological Review, 97 (2), 309–312,

Colonius, H., & Diederich, A. (2020). Formal models and quantitative measures of multisensory integration: a selective overview. European Journal of Neuroscience, 51, 1161–1178,

Colonius, H., & Diederich, A. (2023). Modeling response inhibition in the stop signal task. F.G. Ashby, H. Colonius, & Dzh (Eds.), New handbook of mathematical psychology volume 3 (Vol. 3, pp. 311–356). Cambridge University Press.

Colonius, H., Jahansa, P., Joe, H., Diederich, A. (2024). Towards dependent race models for the stop-signal paradigm: The copula approach. Computational Brain & Behavior, 7 (2), 255–267,

Colonius, H., Özyurt, J., Arndt, P.A. (2001). Countermanding saccades with auditory stop signals: testing the race model. Vision Research, 41 (15), 1951–1968,

Czado, C. (2019). Analyzing dependent data with vine copulas: A practical guide with R. Springer International Publishing.

Durante, F., & Sempi, C. (2016). Principles of copula theory. Chapman and Hall/CRC.

Joe, H. (2014). Dependence modeling with copulas. CRC press.

Logan, G.D., & Cowan, W.B. (1984). On the ability to inhibit thought and action: A theory of an act of control. Psychological review, 91 (3), 295,

Luce, R.D. (1986). Response times: Their role in inferring elementary mental organization (1st ed.). New York, NY 10016: Oxford University Press.

Matzke, D., Dolan, C., Logan, G., Brown, S., Wagenmakers, E.-J. (2013). Bayesian parametric estimation of stop-signal reaction time distributions. Journal of Experimental Psychology: General, 142 (4), 1047–1073,

Matzke, D., Verbruggen, F., Logan, G. (2018). The stop-signal paradigm. J.T. Wixted (Ed.), Stevens’ handbook of experimental psychology and cognitive neuroscience: Methodology (4th ed., Vol. 4, pp. 383–427). John Wiley & Sons.

Nair, J., Wierman, A., Zwart, B. (2022). The fundamentals of heavy tails: Properties, emergence, and estimation (Vol. 53). Cambridge University Press.

Nelsen, R.B. (2006). An introduction to copulas. Springer.

Raab, D. (1962). Statistical facilitation of simple reaction time. Transactions of the New York Academy of Sciences, 24, 574–590,

Sklar, A. (1959). Fonctions de répartition à n dimensions et leurs marges. Publications de l’Institut de Statistique de l’Université de Paris, 8, 229–231,

Townsend, J., & Ashby, F.G. (1983). The stochastic modeling of elementary psychological processes. Cambridge CB2 lRP, UK: Cambridge University Press.

Townsend, J., & Wenger, M. (2004). A theory of interactive parallel processing: new capacity measures and predictions for a response time inequality series. Psychological Review, 111 (4), 1003–1035,

Verbruggen, F., Aron, A.R., Band, G., Beste, C., Bissett, P.G., Brockett, A.T., … Boehler, C.N. (2019). A consensus guide to capturing the ability to inhibit actions and impulsive behaviors in the stop-signal task. eLife, 8, 10.7554/eLife.46323,

